# Mechanistic modeling of amyloid oligomer and protofibril formation

**DOI:** 10.1101/2023.09.02.556028

**Authors:** Keisuke Yuzu, Hiroshi Imamura, Takuro Nozaki, Yuki Fujii, Shaymaa Mohamed Mohamed Badawy, Ken Morishima, Aya Okuda, Rintaro Inoue, Masaaki Sugiyama, Eri Chatani

## Abstract

Early phase of amyloid formation, where prefibrillar aggregates such as oligomers and protofibrils are often observed, is crucial for elucidating pathogenesis. However, since oligomers and protofibrils form transiently and heterogeneously, the detailed mechanisms of their formation remain unclear. Here, we have investigated the early aggregation process of bovine and human insulin by static and dynamic light scattering in combination with thioflavin T (ThT) fluorescence and Fourier transform infrared (FTIR) spectroscopy. The time dependence of light scattering has revealed that oligomers and protofibrils form in bovine insulin, in contrast to no significant aggregation in human insulin. By focusing on bovine insulin for kinetic analysis, it has been revealed that the protofibril formation process was divided into two steps with reference to fractal dimension. When modeled the experimental data of static and dynamic light scattering based on the Smoluchowski aggregation kinetics with fractal aggregation and end-to-end association, we found the initial formation of spherical oligomers and their subsequent uniaxial docking. Furthermore, the analysis of temperature and salt concentration dependence revealed that the end-to-end association is the rate-limiting step, where structure organization occurred with dehydration. The established model for protofibril formation where oligomers are incorporated as a precursor provides insight into the molecular mechanism how protein molecules assemble during the early stage of amyloid formation.

**Significance:** Amyloid oligomers and protofibrils have attracted attention as critical causes of neurodegenerative diseases; however, detailed formation processes of these aggregates have been poorly understood. In this study, we established a mechanistic model of oligomer and protofibril formation of bovine insulin based on Smoluchowski aggregation kinetics in terms of static and dynamic light scattering. It has been demonstrated that early aggregation proceeds by initial fractal-like aggregation to form oligomers, and subsequent their end-to-end docking to form protofibrils. The latter step is a rate-limiting step, where structural organization occurs with dehydration. The established model is expected to broadly applicable to a variety of proteins, and thus will provides valuable insights for accelerating therapeutic development and anti-neurodegenerative drug design.

## Introduction

Amyloid fibrils are fibrillar protein aggregates, the deposition of which is a hallmark of pathology in amyloidoses and neurodegenerative diseases (1). They commonly consist of an ordered cross-β structure and needle-like morphology (2-4), and their formation follows a nucleation-dependent polymerization mechanism. The simplest mechanism of nucleation assumes one-step, where no thermodynamically stabilized assembly species populate (5, 6). Albeit this theoretical assumption, however, a variety of early aggregates are often identified in various proteins. While the size and structure of early aggregates vary depending on proteins and reaction conditions, annular oligomers and rod-like protofibrils are experimentally captured before nucleation (7-9). The oligomers and protofibrils are smaller in size and have more hydrophobic surface properties compared to those of mature amyloid fibrils. Therefore, it has been suggested that they are prone to induce neurotoxicity through the damage of plasma membrane and the binding to signaling receptors (10). The deep involvement of these aggregates is prominent in particular in the pathology of Alzheimer disease (AD), where oligomeric aggregates of amyloid-β have been implicated as a primary agent in neuronal dysfunction (11, 12). Indeed, it is recently worth noting that targeting these species is leading to breakthroughs in immunotherapy of AD (13, 14).

In this context, elucidating the mechanism of protein assembly at the early stages of amyloid formation has become an important issue from the perspective of not only protein chemistry but also pathology and medicine, and emphasis has been placed on mechanistic studies at the protein molecular level. Nevertheless, details of the formation of oligomers and protofibrils remain elusive because these species are often heterogeneous, and furthermore, their population is small. In addition, some species act as on-pathway intermediates (15-18), while others do off-pathway (19, 20), and these features taken together make the time-dependent evolution of early aggregates difficult to be identified. Although many studies have employed schemes where protofibrils are formed via oligomers, it is often speculative and not supported by experimental evidence sufficiently. Quantitative elucidation of molecular mechanism of the early stages of amyloid formation is required for obtaining clues for comprehensive understanding of protein assembly during the formation of oligomers and protofibrils.

To address a more exact mechanism for the formation of amyloid oligomers and protofibrils, we have investigated the time-dependent process of early aggregation using insulin as a representative amyloidogenic model mainly based on static and dynamic light scattering, and in addition, thioflavin T (ThT) fluorescence and Fourier transform infrared (FTIR) spectroscopy. Insulin, a hormone protein essential for glucose metabolism, is one of the best model proteins for studying amyloid formation mechanism as it shows high kinetic quantitativity in amyloid fibril formation in vitro (21-23). Insulin is a 51-residue protein consisting of A-chain with 21 residues and B-chain with 30 residues that are connected by two interchain disulfide bonds. We previously identified in the fibrillation process of insulin from bovine under high NaCl concentration and at high temperature, where protofibril-like aggregates populate transiently before the formation of amyloid fibrils (16, 24). Given that the transformation of protofibrils to amyloid fibrils is enhanced with supply of heat as external energy, they are suggested to serve as an on-pathway intermediate of amyloid fibrils.

By focusing on the protofibril formation of bovine insulin, an early-stage aggregation model has been constructed in this study based on the time course of association number and hydrodynamic radius estimated from the light scattering data. The early aggregation of bovine insulin exhibited size development while maintaining a monomodal distribution, which has enabled the model construction based on Smoluchowski’s scheme applied for colloidal aggregation (25, 26). The whole reaction was divided into two types of assembly, i.e., an isotropic protein assembly to form oligomers, which is then followed by an anisotropic end-to-end association to form protofibrils. Interestingly, insulin from human, which has been investigated in parallel with bovine, did not exhibit early aggregation, from which the sensitivity of early aggregation to amino-acid sequence differences has also been suggested.

## Materials and Methods

### Materials

Bovine insulin was purchased from Sigma-Aldrich Co. Ltd. (St. louis, MO). Recombinant human insulin was purchased form Wako Pure Chemical Industries, Ltd. (Osaka, Japan). ThT was purchased from Wako Pure Chemical Industries, Ltd. HCl solution and NaCl were purchased from Nacalai Tesque, Inc. (Kyoto, Japan).

### Formation of amyloid fibrils

Bovine and human insulin were dissolved in 30 mM HCl and whose concentrations were determined using an absorption coefficient of 1.08 (mg/mL)^-1^ cm^-1^ at 276 nm (27). The spontaneous fibrillation of bovine and human insulin was performed by heating under acidic conditions; various concentrations of insulin from 1.0 to 20 mg/mL in 25 mM HCl containing NaCl at different concentrations ranging from 0 to 1.0 M, were heated at 75 °C with a thermo-regulated heat block (Dry Thermo Unit DTU-1B; TAITEC, Nagoya, Japan).

### ThT fluorescence assay

To monitor the formation of amyloid fibrils, ThT fluorescence assay was performed. At different time points, a 4.5 μL aliquot of the sample solution was mixed with 1.5 mL of 5 μM ThT in 50 mM Gly-NaOH buffer (pH 8.5), and the fluorescence intensity at 485 nm was measured using an excitation wavelength of 445 nm with a RF-5300 PC spectrofluorometer (Shimadzu Corporation, Kyoto, Japan).

### FTIR absorption spectroscopy

FTIR spectra were measured with a Jasco FT/IR-4700ST spectrometer equipped with a DLATGS detector (Jasco, Tokyo, Japan). Prior to the sample preparation, native insulin was preincubated in 25 mM DCl and at 37 °C for 24 h to complete the H/D exchange of amide protons, and 5.0 mg/mL of insulin solution dissolved in 25 mM DCl containing 0.5 M NaCl was heated at 75 °C. At different time points, an approximately 50 μL aliquot of the sample, which was air-cooled to room temperature, was sealed with a CaF_2_ window cell and a 50 μm polytetrafluoroethylene spacer embedded inside the water-circulating system connected to a thermoregulated water bath. FTIR spectra were then measured at 25 °C by collecting 256 interferograms at a resolution of 2 cm^−1^. The obtained spectra exhibited the amide Ⅰ’ bands, which was slightly shifted to lower frequencies than the amide I bands due to the substitution of amide protons by deuterium.

### Static and dynamic light scattering

To probe the evolution of protein aggregates, static and dynamic light scattering measurements were performed using a Zetasizer Nano S (Malvern Instruments Ltd., Worcestershire, U.K.) with *θ* = 173° and *λ* = 632.8 nm. The heat-induced aggregation of insulin was initiated instantaneously by mixing an insulin solution with preheated other components, and the sample solution was immediately transferred to a preheated quartz microcuvette embedded inside the thermo-regulating system. The temperature jump of the sample solution to the reaction temperature was completed within approximately 60 sec. Each light scattering data during the time course measurement was collected using an acquisition time of 5 sec.

The normalized time auto correlation function of the scattering intensity for a given correlation time *τ, g*^(2)^(*τ*), was defined by

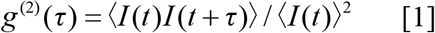

where *I*(*t*) and *I*(*t + τ*) are the scattering intensities at time *t* and *t* + *τ*, respectively. The angle brackets indicate averaging over time. *g*^(2)^(*τ*) is related to the normalized electric field-field time autocorrelation function, *g*^(1)^(*τ*), via the following equation:

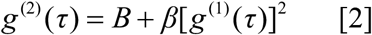

where *B* is the long-time value of *g*^(2)^(*τ*) (*B* ∼1) and *β* is an instrument parameter. The moment method by Frisken (28), which is a reformulated version of the method of cumulants gives

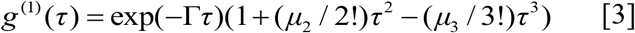

where Γ is the mean decay rate, and *μ*_m_ are the *m*th moments about the mean, which correspond directly to the *m*th cumulants in the method for cumulants (28). The functions of eqs. 2 and 3 with terms up to the second moment about the mean, *μ*_2_, were used for the fitting analysis. *Γ* is given by

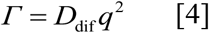

where *D*_dif_ is the diffusion coefficient of the particles. *q* is the magnitude of the scattering vector defined as

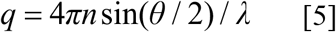

where *n* is the refractive index of the solvent, *θ* is the scattering angle, and *λ* is the wavelength of the light. The diffusion coefficient, *D*_dif_, is rationalized to the hydrodynamic radius of the particle, *R*_H_, via the Stokes-Einstein relation:

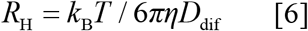

where *k*_B_ is the Boltzmann constant, *T* is the thermodynamic temperature, and *η* is the viscosity of the solvent. Size distribution weighted by light scattering intensity was also obtained by the non-negative least-squares (NNLS), a regulation method for numerical inversion of the Laplace transform implemented in Zetasizer software version 7.13 (Malvern Instruments Ltd.).

The excess static light scattering from the protein particles, *I*_ex_, is determined by <*I*(*t*)>_sol_ – <*I*(*t*)>_solv_, where subscripts “sol” and “solv” denote a protein solution and a solvent, respectively. The relative excess light scattering intensity, *I*/*I*_0_, is calculated by

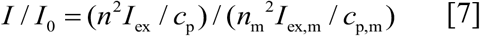

where *c*_p_, *I*_ex,m_, *c*_p,m_, and *n*_m_ are the mass concentration of the particle, the excess light scattering intensity from the primary (monomeric) particle, its mass concentration, and the refractive index of the dissolving solvent, respectively. For Rayleigh scattering, the excess light scattering intensity is rationalized by the Rayleigh ratio, *R*(*θ*):

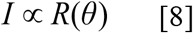

*R*(*θ*) is expressed as (29)

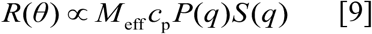

where *P*(*q*) is the form factor, i.e., the intraparticle or the intracluster interference, and *S*(*q*) is the structure factor, i.e., the interparticle or the intercluster interference. *M*_eff_ is the effective molecular weight of the aggregate, given as

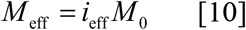

where *i*_eff_ is the effective number of the primary (monomeric) particles in the aggregate and *M*_0_ is the molecular weight of the monomer. Here, we assumed *P*(*q*) = 1, which is a reasonable approximation when the size of a particle is smaller than *q*^-1^, i.e., *qR*_H_ < 1 (29, 30). *q*^-1^ was 38 nm in the present experimental setup. The present experimental conditions under which the protein was sufficiently diluted and satisfies *S*(*q*) = 1., where the interparticle effect on the scattering is negligibly small. Accordingly, the excess light scattering intensity is proportional to the molecular weight: eqs. 7-10 give

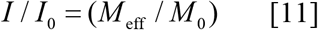

## Results

### Amyloid fibril formation of bovine and human insulin

We first characterized the fibrillation process of bovine and human insulin, the amino acid residues of which differ by only three out of 51 residues (Figure 1A), by using ThT fluorescence assay. In our previous study, the formation of protofibrils showing high ThT fluorescence intensity has been identified in bovine insulin at high NaCl concentration and high temperature (16, 24). Indeed, an explosive increase and subsequent decrease in ThT fluorescence were observed when bovine insulin solution at 5.0 mg/mL was heated in the presence of various concentrations of NaCl (Figure 1B). The protofibrils accumulated in a larger amount and with a longer lifetime under 0.5 M NaCl than 0.1 and 0.2 M NaCl conditions, suggesting that the protofibrils were more favorably produced at higher NaCl concentrations. Unexpectedly, a sigmoidal pattern with a lag time in ThT fluorescence, which is attributed to a typical nucleation-dependent polymerization mechanism, was observed in human insulin when measured at the same protein and NaCl concentrations (Figure 1C). These results indicate that the fibrillation process via protofibrils is intrinsic to bovine insulin, albeit the amino acid sequences between bovine and human insulin differ only slightly. The bovine insulin-specific formation of protofibrils was also identified by the construction of phase diagrams (Figure S1) and small-angle X-ray scattering (SAXS) (Figure S2).

**Figure 1.**
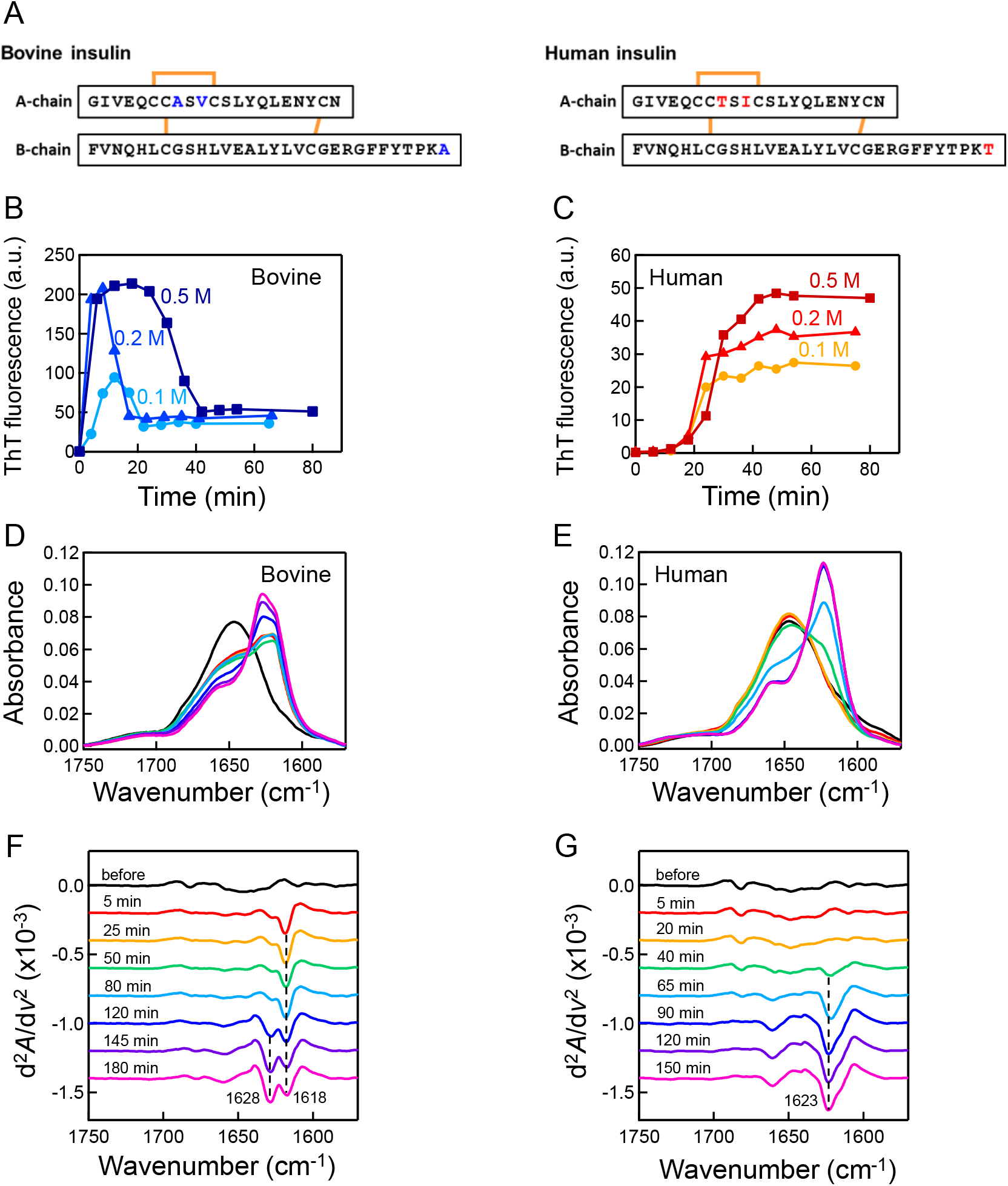
Amyloid fibril formation pathway of bovine and human insulin. (A) Amino acid sequences of bovine and human insulin. The red and blue characters represent different residues between bovine and human insulin, respectively (B, C) Time courses of ThT fluorescence intensity in amyloid fibril formation of bovine (B) and human insulin (C). Amyloid fibril formation was induced by heating at 75 °C under the conditions of 5.0 mg/mL insulin and 0.1-0.5 M NaCl. (D, E) Time dependence of FTIR spectra at the amide I’ region in amyloid fibril formation of bovine (D) and human insulin (E). Amyloid fibril formation was performed under the conditions of 5.0 mg/mL insulin and 0.5 M NaCl at 75 °C. At different time points, an aliquot of the reaction solution, which was air-cooled to room temperature, was sealed with a cell and measured at 25 °C. Spectra were normalized so that the integrated intensity of the amide I′ band ranging from 1580 to 1750 cm^−1^ was set to be equal. (F, G) Second derivative FTIR spectra of bovine (F) and human insulin (G). The dashed black lines indicate the positions of 1618 cm^-1^ and 1628 cm^-1^ for bovine insulin and 1623 cm^-1^ for human insulin.

We also performed time-resolved FTIR measurement to verify the difference in the fibrillation process between bovine and human insulin, where deuterated conditions were used to observe the amide region without any interference of water absorption. The basic ThT fluorescence pattern of time course was conserved in both insulin species even in deuterated solution, although the fibrillation rate became slower (Figure S3). In both types of insulin, a sharp β-sheet peak was observed at around 1620-1630 cm^-1^ in the amide Ⅰ’ region as fibrillation progressed (Figure 1D, E). As observed in the second derivative spectra, the β-sheet peak at 1618 cm^-1^ appeared abruptly and another β-sheet peak at 1628 cm^-1^ developed thereafter for bovine insulin (Figure 1F), supporting the appearance of protofibrils prior to the formation of mature fibrils (16, 24). In contrast, the spectral shape was unchanged significantly during the lag time and only a single β-sheet peak at 1623 cm^-1^ developed cooperatively with ThT increase for human insulin (Figure 1G), supporting the formation of mature fibrils without any prefibrillar aggregates. Furthermore, the differences in β-sheet peak of bovine and human insulin suggested their structural difference consistent with our previous study (31).

### Monitoring bovine insulin-specific early aggregation by dynamic light scattering

We next carried out dynamic light scattering measurements to track the formation of prefibrillar aggregates of bovine insulin in a comparative manner with human insulin. When the time-dependent evolution was monitored after temperature jump, the decay of autocorrelations, *g*^(2)^(*τ*)-1, of both bovine and human insulin exhibited time-dependent increase, indicating that the size of protein aggregates evolved with the progression of time (Figure 2A, B). Interestingly, the autocorrelations of bovine insulin developed rapidly maintaining a single exponential pattern, and the data analysis by NNLS method exhibited a rapid growth of aggregates with a monomodal distribution (Figure 2C). On the other hand, the autocorrelations showed a slow evolution in human insulin. Although they apparently showed multiple exponential patterns, the data analysis by the NNLS method clarified that this evolution is attributed to the formation of a small fraction of large aggregates. The distribution of the aggregates became almost negligible when considering it as a volume fraction, and therefore it was concluded that early aggregation did not proceed in human insulin (Figure 2D).

**Figure 2.**
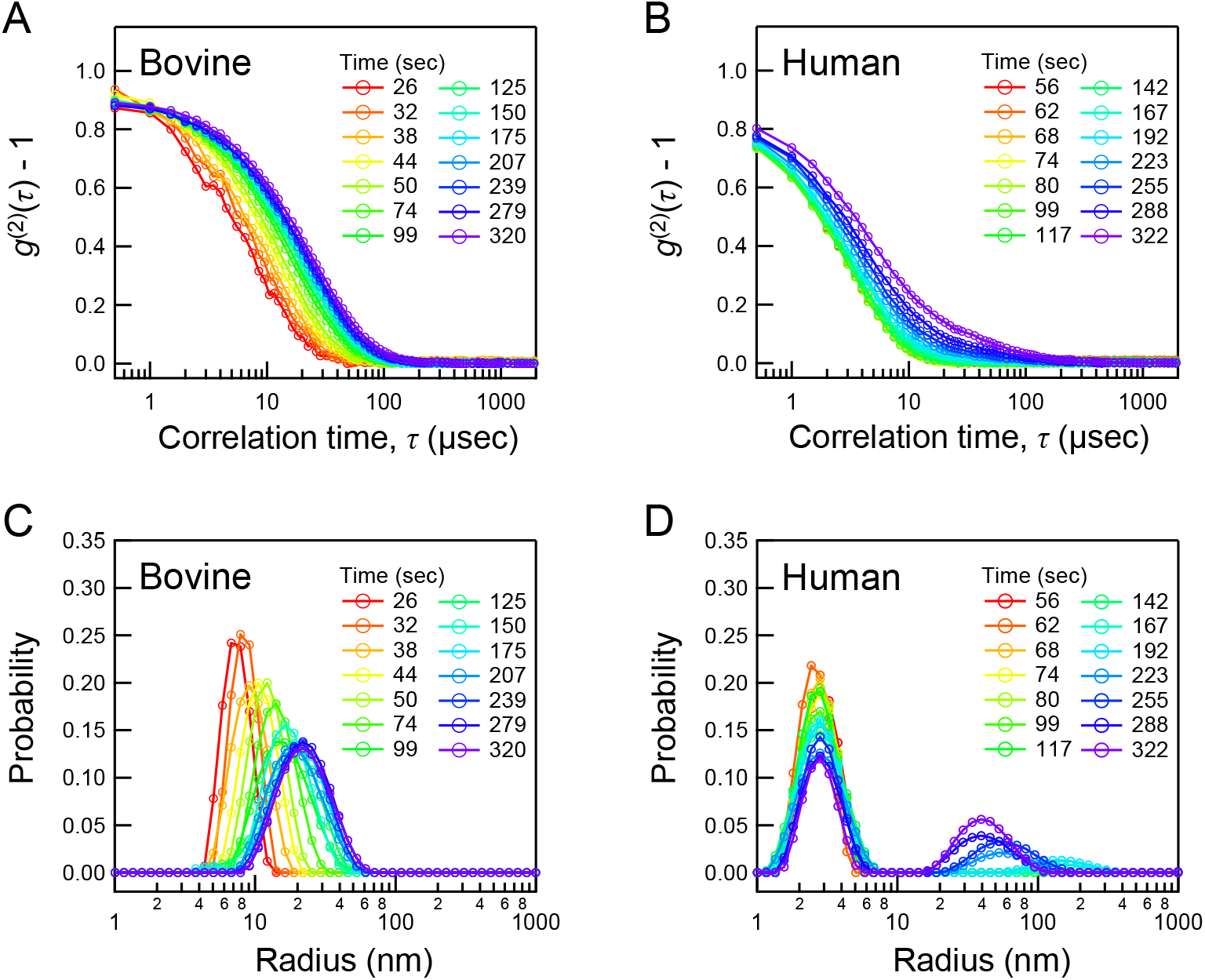
Time dependence of dynamic light scattering during the early aggregation phase of amyloid fibril formation. The heat-induced aggregation of insulin was performed under the conditions of 5.0 mg/mL insulin and 0.5 M NaCl, and monitored for approximately 5 min after initiation of the reaction by temperature jump to 75 °C. (A, B) Time dependence of the normalized time autocorrelation functions of bovine (A) and human insulin (B). (C, D) Intensity-weighted distribution of the size of bovine (C) and human insulin (D) determined by NNLS analysis on the autocorrelations.

These results demonstrate that bovine insulin forms protofibrils in the early stage of fibrillation, in contrast to typical one-step nucleation without any thermodynamically metastable aggregates. Furthermore, the aggregates kept monomodal and relatively narrow size distribution during the measurement time period in Figure 2C, which has indicated that the data are suitable for more quantitative analyses. The gradual and monomodal size development was specific to bovine insulin, which was also supported by analytical ultracentrifugation measurements (Figure S4). We hereafter focused on the time-dependent process of early aggregation of bovine insulin in the current reaction conditions using dynamic and in addition, static light scattering.

### Estimating the number of phases in the aggregation of bovine insulin

We analyzed data from concurrent monitoring of dynamic and excess static light scattering to grasp the framework of the aggregation kinetics of bovine insulin. The time dependence of hydrodynamic radius *R*_H_, a parameter determined from the autocorrelation function of dynamic light scattering according to eqs. 4-6, showed explosive growth and subsequent gradual development (Figure 3A), indicating that the early aggregation proceeds in two steps. The relative light scattering intensity, *I*/*I*_0_ obtained from static light scattering according to eqs. 7-10, which corresponds to effective association number of protein particles in an aggregate, also showed a similar pattern of size evolution to that of *R*_H_ (Figure 3B). In the first step, as the protein concentration decreased, a slight delay became more prominent by approximately 30 sec after initiating the reaction (Figure 3C). This result suggests that the first step involves a lag phase before the explosive growth.

**Figure 3.**
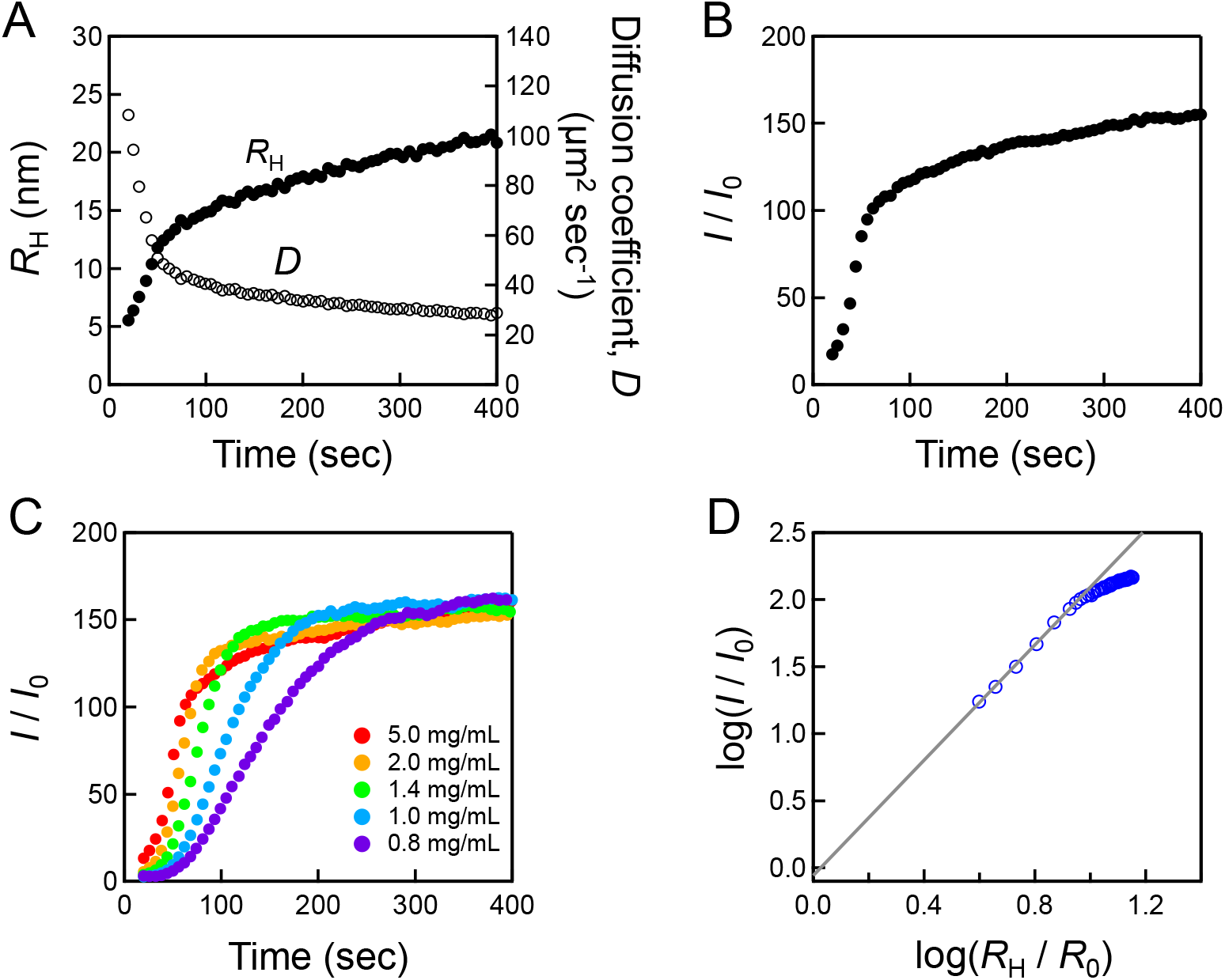
Aggregation kinetics during the early aggregation phase of bovine insulin. (A) Time dependence of the diffusion coefficient and the hydrodynamic radius. (B) Time dependence of the relative excess static light scattering intensity. (C) Protein concentration dependence on the early aggregation kinetics. Time dependence of the relative excess static light scattering intensity was measured in the range of 0.8-5.0 mg/mL. (D) Power-law relationship between excess static light scattering intensity and hydrodynamic radius. We applied the fitting analysis with eq. 17 to the data region up to approximately 60 sec after initiating the reaction, which satisfies *R*_H_ < 38 nm (*qR* < 1). The solid line represents the fitting line with a slope of 2.15 and an intercept of -0.06 ± 0.03.

To further explore the character of aggregates formed in each step, we examined the time dependence of fractal dimension, *D*_f_, which reflects the packing of primary particles in the aggregate, i.e., compactness or morphology of the aggregates (32). *D*_f_ obeys the following relationship: (33, 34)

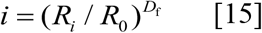

where *i* is the number of a primary particle in a cluster, *R*_i_ is the radius of the cluster of an *i*-mer, and *R*_0_ is the radius of a primary particle. Here, the primary particle is regarded as a monomer, and the hydrodynamic radius *R*_H_ corresponds to *R*_i_ in this equation (26). According to eqs. 10 and 15, the effective number of primary particles, *i*_eff_, in a cluster is written as

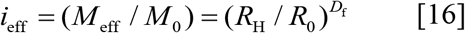

Thus, the power-law relationship between the relative excess light scattering intensity and relative hydrodynamic radius is obtained as

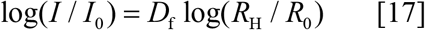

Here, the fractal dimension obtained from the slope of log(*I*/*I*_0_) against log(*R*_H_/*R*_0_) was *D*_f_ = 2.15 ± 0.04 in the range up to approximately 60 sec, corresponding to the boundary of the first and second steps. This value is closer to that for reaction-limited cluster aggregation (*D*_f_ = 2.05) (34, 35) than that for diffusion-limited cluster aggregation (*D*_f_ = 1.75) (36).

However, the time dependence of log(*I*/*I*_0_) against log(*R*_H_/*R*_0_) gradually deviated from the original slope in the second step (Figure 3D). In light of our previous finding that bovine insulin forms protofibril-like prefibrillar aggregates before the formation of amyloid fibrils (24), it was presumed that the association mode of the clusters changed from spherical to rod-like. Based on this result, we presumed that fractal aggregation-like spherical association proceeds in the first step, which is followed by anisotropic association towards a rod in the second step.

### Establishment of an aggregation model of bovine insulin

Based on the above-mentioned characterization that the aggregation of bovine insulin consists of two different association modes, the aggregation process of bovine insulin was modeled based on Smoluchowski aggregation kinetics. In this theory, aggregation is formulated by two diffusing particles colliding and irreversibly coagulating with a given probability (25), as represented by the following equation of the time dependence of the concentration (the fraction or the population) of all *i*-mers:

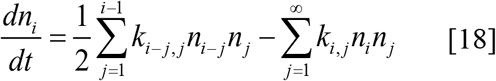

where *n*_*i*_ is the concentration of *i*-mer. The concentration is normalized to the initial primary particle concentration (∑_*i*_*in*_*i*_ = 1). *t* is time, which was set to zero when heating of protein solution was initiated. *k*_*i,j*_ is the rate constant at which an *i*-mer and a *j*-mer collide by diffusion per unit time. *k*_*i,j*_ is given as

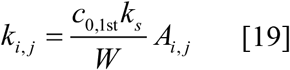

where *c*_0,1st_ is the initial concentration of the primary particle (the initial number density of the monomer) in the first step, i.e., *c*_0,1st_ = 5.25×10^23^, *k*_s_ is the Smoluchowski rate constant (*k*_s_ = 8*k*_B_*T*/3*η*), and *W* is Fuchs stability ratio, which describes the interparticle interaction between two particles (37). *A*_*i,j*_ is the correlation term of the probability that an *i*-mer and a *j*-mer collide by diffusion per unit time. For fractal-like aggregation of spherical aggregates in the first step (Figure 4A), *A*_*i,j*_ is described as:

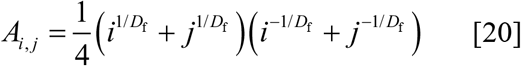

which is a modified description with the concept of the fractal to the original formula by Smoluchowki (34, 38, 39). Here, two different *W* values were adopted to reproduce the lag phase and the subsequent rapid aggregation, respectively, in the first step as observed in Figure 3C. For end-to-end aggregation of rod-like aggregates in the second step (Figure 4B), *A*_*i,j*_ in eq. 19 is replaced by another correlation term, *B*_*i’,j’*_, which describes the probability that an *i*’-mer and a *j*’-mer collide by diffusion per unit time (40):

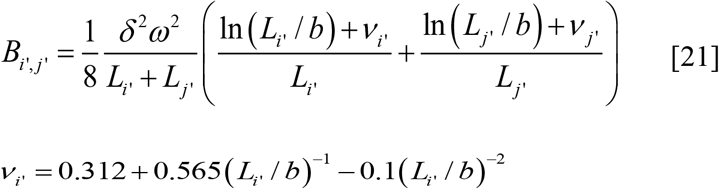

**Figure 4.**
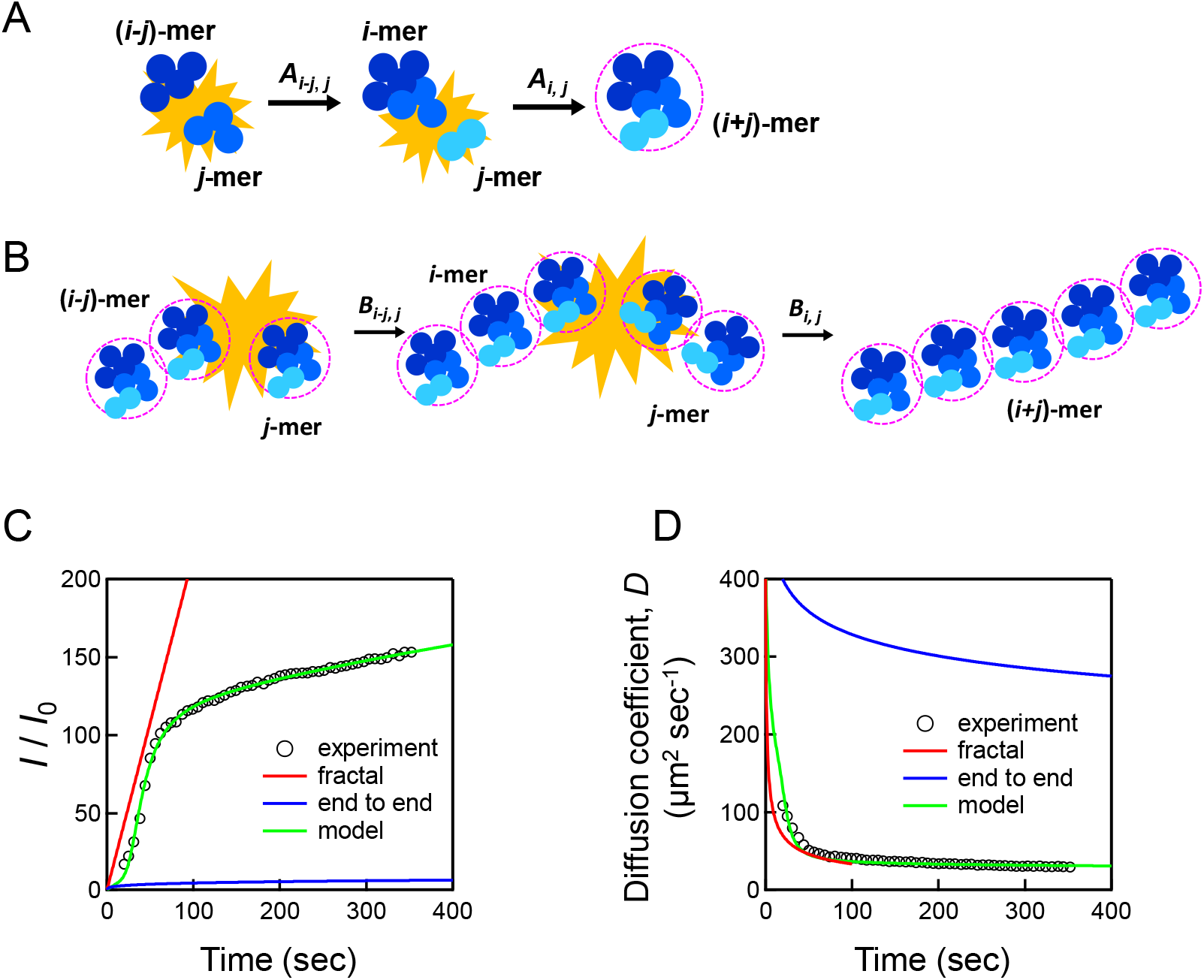
Modeling the early aggregation of bovine insulin based on Smoluchowski aggregation kinetics. (A, B) Schematic illustration of the present aggregation model for fractal (A) and subsequent end-to-end aggregation (B). (C, D) Time dependence of the relative excess static light scattering intensity (C) and the diffusion coefficient (D) simulated by the Smoluchowski aggregation equation. The black circles indicate experimental data. The red, blue, and green lines indicate the curves simulated by fractal aggregation, end-to-end aggregation, and the present multistep model, respectively.

Here, it should be noted that *i*’ and *j*’ were defined as the association number of spherical aggregates formed in the first step. Therefore, *k*_*i’,j’*_ in the second step is given as,

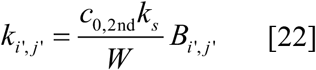

Where *L*_*i’*_ is the length of rod of an *i’*-mer. *δ* and *ω* are the minimum allowed distance and angle between two colliding rods, respectively. *δω* was defined as *δω* = *d*(*i’*+*j’*)/2*i’j’* as described previously (41), while in the present work, we considered that *δ* and *ω* change depending on the length of the rod. These distinct aggregation modes were switched at a certain threshold cluster size, and the size of fractal aggregated clusters was set to the threshold before the start of end-to-end aggregation; therefore, the initial concentration of the primary particle in the second step, *c*_0,2nd_, was set to 2.38×10^21^.

The aggregation equations were numerically solved considering the clusters with *i* < 7500, which was enough to conserve the total mass of the system in eq 18 approximating to an infinite system. Based on this, the excess light scattering intensity was calculated according to (39)

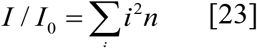

The Smoluchowski equation describes the time dependence of the population of the *i*-mer, *in*_*i*_. Then, the diffusion coefficient of a system composed of an ensemble of aggregates was calculated by

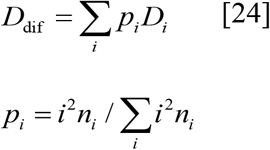

where *D*_*i*_ is the diffusion coefficient of the *i*-mer, and *p*_*i*_ is the normalized intensity-weighted distribution of the *i*-mer.

The theoretical time dependence of *I*/*I*_0_ and diffusion coefficient defined by eqs. 23 and 24 successfully reproduced the experimental data (green lines in Figure 4C, D). Either fractal or end-to-end aggregation model alone could not reproduce the data (red and blue lines, respectively, in Figure 4C, D), verifying that the current aggregation requires the combination of these two aggregation mechanisms. Through fitting the model curve to the experimental plots, the thresholds of the lag phase in the fractal aggregation and of the shift to the end-to-end association mechanism have been predicted to be ∼30 mer (*R*_30_ ∼ 6.6 nm) and ∼150 mer (*R*_150_ ∼ 13.7 nm), respectively. The values of *W* have also been predicted to be 4.6 (±0.1) ×10^7^ (lag phase) and 3.5 (±0.1) ×10^6^ (growth phase) for the first fractal aggregation and 1.7 (±0.1) ×10^9^ for the subsequent end-to-end association. These values are clearly larger than 1, indicating that the early aggregation of bovine insulin progresses in a reaction-limited manner in all steps.

### Effect of temperature and salt concentration on the early aggregation

We further measured the dependence of temperature and NaCl concentration to examine detailed characteristics of the aggregation. When *I*/*I*_0_ was monitored in the range of 70-80 °C, the time course showed significant acceleration with the increase of temperature (Figure 5A). To examine activation enthalpy, Δ*H\**, in each aggregation step, we analyzed an Arrhenius plot of an effective rate constant of aggregation, *k*_eff_ *= k*_s_/*W*, where the specific probability of the reaction is taken into account by combining *W* with *k*_s_. The values of *W* and *k*_eff_ obtained by fitting analysis are shown in Table S1. Δ*H\** is expressed as (38)

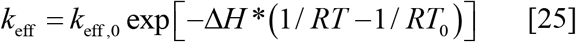

where *k*_eff,0_ is *k*_eff_ at the reference temperature, *T*_0_, and *R* is the gas constant. Δ*H\** was obtained from the slope, the value of which were 133 ± 8 kJ/mol (lag phase) and -14 ± 14 kJ/mol (growth phase) for the initial fractal aggregation, and 177 ± 24 kJ/mol for the end-to-end association (Figure 5B). It has been revealed that the pathway consists of two dominant energy barriers for the progression of aggregation, and the latter end-to-end association acts as a rate-limiting step in the whole reaction.

**Figure 5.**
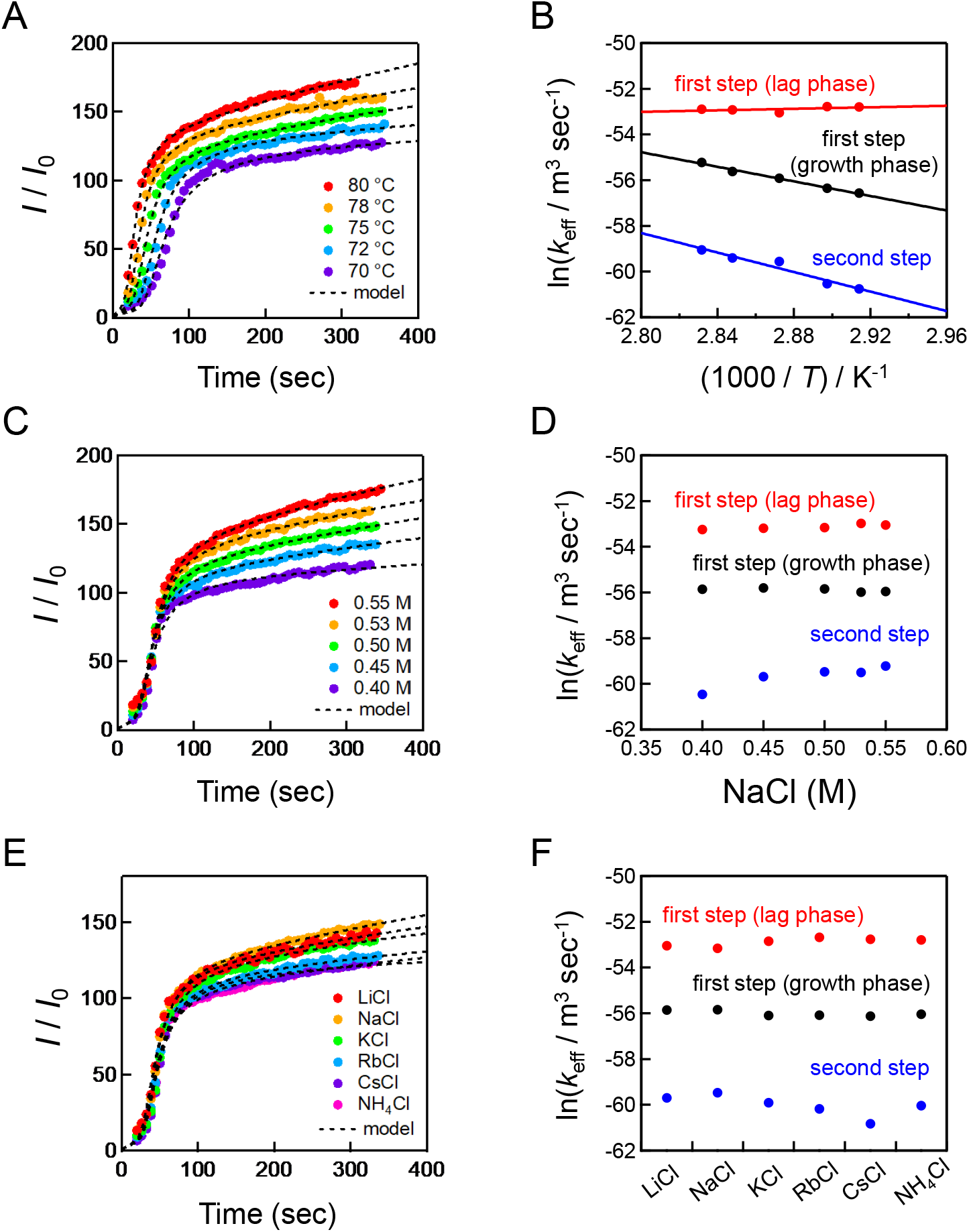
(A) Temperature dependence on the early aggregation of bovine insulin in the range of 70 to 80 °C. Time dependence of relative excess static light scattering intensity is shown. (B) Temperature dependence on the effective aggregation rate constant, *k*_eff_, for first (lag and growth periods) and second steps. *k*_eff_ was determined by the relative excess light scattering intensity, *I/I*_0_. The solid lines indicate the fitting line by eq. 25 in the range between 70 to 80 °C. (C) Salt concentration dependence on the early aggregation of bovine insulin. Time dependence of relative excess static light scattering intensity in the range of 0.4 to 0.55 M NaCl is shown. (D) Salt concentration dependence on the effective aggregation rate constant for first (lag and growth periods) and second steps. (E) Hofmeister salt dependence on the early aggregation of bovine insulin. Time dependence of relative excess static light scattering intensity of various salts with different cations is shown. (F) Hofmeister salt dependence on the effective aggregation rate constant for first (lag and growth periods) and second steps.

The concentration dependence of NaCl was also monitored in the range of 0.40-0.55 M. When *I*/*I*_0_ was monitored, its time evolution showed selective acceleration in the end-to-end association (Figure 5C), which has been clearly demonstrated by *k*_eff_ determined by curve fitting (Figure 5D and Table S2). Considering that the NaCl concentrations used for the analysis are high enough to electrostatically shield protein molecules (Figure S5), it is expected that the salt-induced acceleration is related to the hydration state, the properties of which tend to be modulated by ionic interactions with protein molecules and water. In agreement with this idea, *k*_eff_ of the end-to-end association showed relationship to Hofmeister series when NaCl was substituted for other salts with different cations (Figure 5E, F and Table S3).

## Discussion

### Early aggregation process of bovine insulin

In this study, we have achieved to describe the protofibril formation mechanism of bovine insulin based on Smoluchowski aggregation kinetics. Protofibrils have attracted attention as a precursor of amyloid fibrils, and recently, they have also been focused on as pathologically important aggregates that exhibit cytotoxicity along with oligomers (12). However, the detailed process of protofibril formation has been poorly tracked, often due to the low and heterogeneous distribution of aggregates in the early stage. As a breakthrough for the difficulty, we have found an almost monodisperse development of early aggregates in bovine insulin (Figure 2), which has allowed for the establishment of a mechanistic model of oligomer and protofibril formation.

The Smoluchowski aggregation equation or population balance equation is an attractive basis for analyzing protein aggregation because of its simple assumption of irreversible colliding of two particles. Indeed, it has been applied when analyzing aggregation processes of many peptides and proteins (26, 38-40, 42-47). The examples include studies on amyloid fibril formation (42-45, 47); however, they mainly describe the elongation process of amyloid fibril formation by applying only one type of aggregation mode. Here, we have revealed that the early aggregation of bovine insulin could be divided into two steps based on changes in fractal dimension, and successfully reproduced the experimental data by assuming isotropic fractal aggregation, followed by anisotropic end-to-end elongation. Recently, the development of AmyloFit has greatly advanced our understanding of microscopic steps during amyloid fibril formation, which has been extended to kinetic modeling of oligomer dynamics for therapeutic strategies for their inhibition (48, 49). At present, however, it is limited to revealing the distribution of oligomers with a fixed association number. The schematic model presented in this study is based on the in molecular weight and diffusion coefficient of aggregates, and has seamlessly provided time dependence of aggregates not only in the size but also in the shape.

Figure 6 shows the scheme of oligomer and protofibril formation of bovine insulin revealed in this study. Fractal-like spherical aggregation proceeds in the first step, and their end-to-end association initiates in the second step to form rod-like aggregates when the spherical clusters reach a certain size. This scheme shows that oligomer and protofibril formation essentially progresses by a mechanism close to colloidal aggregation, distinct from the nucleation-dependent growth of amyloid fibrils. The switching of aggregation mode occurs after the size of spherical clusters develops to ∼13.7 nm in hydrodynamic radius (Figure 4). Given that the first step corresponds to oligomer formation, the present scheme describes the protofibril formation mechanism using oligomers as building blocks. This two-step scheme has not been considered in our previous tracking of protofibril formation of bovine insulin by SAXS (24), just by approximating aggregate shape to an elliptical cylinder over the whole measurement time range. However, this analysis also showed initial thickening of the elliptical cylindrical aggregates prior to subsequent elongation, presumably corresponding to the initial formation of spherical clusters.

**Figure 6.**
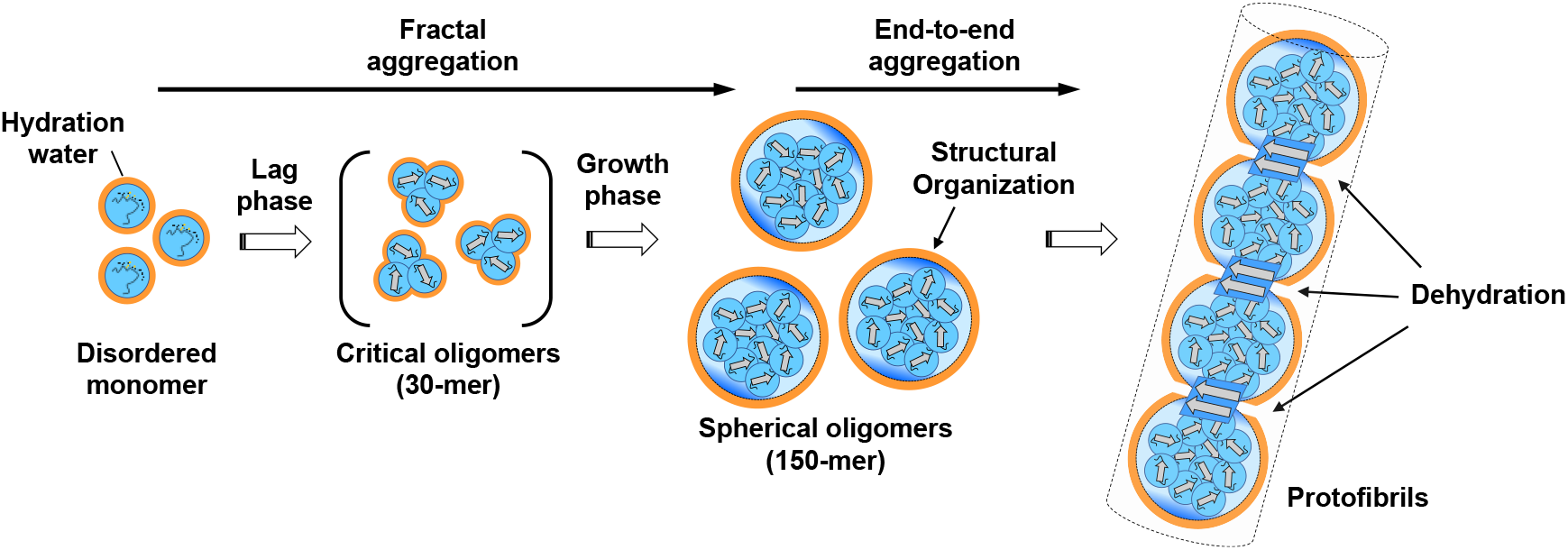
Schematic illustration representing the protofibril formation mechanism of bovine insulin.

### Proposed mechanism of aggregation scheme switching

The proposed scheme involves switching of the two aggregation schemes at a threshold of cluster size. Since the formation of spherical and rod-like aggregates obey different correlation terms, *A*_*ij*_ and *B*_*ij*_, respectively, their formation pathways are theoretically different. However, the present observation suggests that they approach each other as aggregation proceeds, making it possible to switch at the point of closest proximity. Because protofibrils contain some amount of β-sheet structure (Figure 1D, F), their critical free energy is expected to be higher than that of oligomers resulting in an initial kinetic preference for oligomer formation. As aggregates develop in size, however, protofibrils gain more pronounced stability and eventually exceed the stability of oligomers, which may change the aggregation pathway from oligomers to protofibrils. Since hydrophobic interactions are considered to drive oligomer formation, the growth of oligomers will become unfavorable with a threshold where the number of hydrophobic residues is insufficient to fill the interior of the cluster and some of the hydrophilic residues must be buried; this may determine the timing of the switch from spherical oligomers to anisotropically shaped protofibrils with a smaller relative internal volume.

It is also important to clarify what is the driving force for the end-to-end assembly. Although no structural details were obtained directly in this study, the presence of a lag phase in the oligomer formation (Figure 3C) indicates that oligomers contain some degree of ordered structure. Considering that oligomers serve as building blocks for protofibril growth, they are expected to acquire some specific structure that plays a role as binding sites at two ends. Interestingly, a strong salt-induced enhancement was observed in the second step even at high salt concentrations where the charge shielding is fully completed (Figure 5). In addition, the Hofmeister series (50) was highlighted when analyzing various salts (Figure 5E, F), strongly implicating the involvement of water molecules in the end-to-end association. There are several studies proposing interplays between water and the structural dynamics of proteins, where dehydration of polar groups of proteins has been revealed to undergo in the rate-limiting step of protein folding, protofilament association and elongation of amyloid fibrils (51-53). Given that the hydrogen bonds formed by polar groups provide directional specificity, it is presumed that water molecules, which act as hydrogen bond-forming groups, overcome the high dehydration energy to acquire interactions with orientational specificity important for organizing ordered structure. Likewise, aggregate structure becomes more intensely ordered in the end-to-end association, presumably forming a β-structure, through expulsion of water from polar groups at the interface, by paying the cost of dehydration energy barrier.

### Differences in aggregation scheme between bovine and human insulin

Another notable observation in this work is that human insulin showed the simplest one-step nucleation scheme without any significant formation of oligomers and protofibrils (Figure 1, 2). The amino acid sequence differs between bovine and human insulin only by 3 of 51 residues. The disappearance of early aggregates in human insulin despite such high sequence homology implies that the nucleation mechanism is highly dependent not only on the reaction conditions but also on the amino acid sequence. This is quite in analogy to observations that the ΔE22 and E22G mutant of amyloid β produces markedly larger amounts of oligomers than the wild type (54, 55) and that the E46K and A30P mutants of α-synuclein promote and delay oligomer formation, respectively (56). Both of these are considered to be representative examples in living systems in which even slight residue differences affect aggregation pathway.

In our previous study, we identified structured regions of bovine insulin protofibrils by protease digestion. The result demonstrated that the 8th and 10th positions of the A chain were structured in protofibrils (16). Combining this finding with another observation that porcine insulin, in which only the 30th residue of the B-chain is bovine type, showed a one-step nucleation similar to human insulin (data not shown), it is suggested that two residues of the A-chain contribute to oligomer and protofibril formation. The initiation of oligomer and protofibril formation is expected to be driven by hydrophobic interactions and formation propensity of β structures; however, no clear differences between bovine and human insulin have been so far found even using algorithms (57), implying that very slight sequence characteristics may cause variations in amyloid nucleation scheme.

### Insights into oligomer and protofibril formation in biological system

The present work has shed light on one of the essential schemes of amyloidogenic aggregation, in which oligomers serve as on-pathway intermediates for the formation of protofibrils. This is to our knowledge the first model demonstrating the way of how oligomers are incorporated to form protofibrils based on experimental measurements. Both of the formation of oligomers and protofibrils could be described by simple schemes and the established model has successfully characterized each process. The model analysis of bovine insulin became feasible because only a single pathway was ideally observed, while diverse pathways are predicted to progress simultaneously in most cases of amyloid formation in pathology. Although addressing experimentally the full spectrum of aggregation pathways may not be so straightforward, the current scheme of oligomer and subsequent protofibril formation represent the framework of the pathways of early aggregation. It is also a new discovery that the rate-limiting step of protofibril formation is the association of oligomers where the expulsion of water is involved, indicating that the structural organization progresses considerably during this step. Such a detailed picture provides important insights in devising strategic targeting strategies for antibodies and other anti-aggregating agents. Advances in clarifying aggregation mechanisms at the protein molecular level will pave the way for early treatment of amyloidosis and neurodegenerative diseases.

## References

1. P. Westermark et al., A primer of amyloid nomenclature. Amyloid 14, 179–183 (2007).

2. F. Chiti, C. M. Dobson, Protein misfolding, functional amyloid, and human disease. Annu Rev Biochem 75, 333–366 (2006).

3. D. Eisenberg, M. Jucker, The amyloid state of proteins in human diseases. Cell 148, 1188–1203 (2012).

4. M. G. Iadanza, M. P. Jackson, E. W. Hewitt, N. A. Ranson, S. E. Radford, A new era for understanding amyloid structures and disease. Nat Rev Mol Cell Biol 19, 755–773 (2018).

5. F. Ferrone, Analysis of protein aggregation kinetics. Methods Enzymol 309, 256–274 (1999).

6. P. Arosio, T. P. Knowles, S. Linse, On the lag phase in amyloid fibril formation. Phys Chem Chem Phys 17, 7606–7618 (2015).

7. A. Quist et al., Amyloid ion channels: a common structural link for protein-misfolding disease. Proc Natl Acad Sci U S A 102, 10427–10432 (2005).

8. A. E. Langkilde, B. Vestergaard, Methods for structural characterization of prefibrillar intermediates and amyloid fibrils. FEBS Lett 583, 2600–2609 (2009).

9. E. Chatani, N. Yamamoto, Recent progress on understanding the mechanisms of amyloid nucleation. Biophys Rev 10, 527–534 (2018).

10. T. L. Williams, L. C. Serpell, Membrane and surface interactions of Alzheimer’s Abeta peptide--insights into the mechanism of cytotoxicity. FEBS J 278, 3905–3917 (2011).

11. M. Sakono, T. Zako, Amyloid oligomers: formation and toxicity of Abeta oligomers. FEBS J 277, 1348–1358 (2010).

12. T. Yasumoto et al., High molecular weight amyloid beta(1-42) oligomers induce neurotoxicity via plasma membrane damage. FASEB J 33, 9220–9234 (2019).

13. C. H. van Dyck et al., Lecanemab in Early Alzheimer’s Disease. N Engl J Med 388, 9–21 (2023).

14. E. Karran, B. De Strooper, The amyloid hypothesis in Alzheimer disease: new insights from new therapeutics. Nat Rev Drug Discov 21, 306–318 (2022).

15. C. Bleiholder, N. F. Dupuis, T. Wyttenbach, M. T. Bowers, Ion mobility-mass spectrometry reveals a conformational conversion from random assembly to betasheet in amyloid fibril formation. Nat Chem 3, 172–177 (2011).

16. E. Chatani, H. Imamura, N. Yamamoto, M. Kato, Stepwise organization of the beta-structure identifies key regions essential for the propagation and cytotoxicity of insulin amyloid fibrils. J Biol Chem 289, 10399–10410 (2014).

17. A. Pavlova et al., Protein structural and surface water rearrangement constitute major events in the earliest aggregation stages of tau. Proc Natl Acad Sci U S A 113, E127–136 (2016).

18. N. Yamamoto, S. Tsuhara, A. Tamura, E. Chatani, A specific form of prefibrillar aggregates that functions as a precursor of amyloid nucleation. Sci Rep 8, 62 (2018).

19. T. Miti, M. Mulaj, J. D. Schmit, M. Muschol, Stable, metastable, and kinetically trapped amyloid aggregate phases. Biomacromolecules 16, 326–335 (2015).

20. Y. Cao, J. Adamcik, M. Diener, J. R. Kumita, R. Mezzenga, Different Folding States from the Same Protein Sequence Determine Reversible vs Irreversible Amyloid Fate. J Am Chem Soc 143, 11473–11481 (2021).

21. J. Brange, L. Andersen, E. D. Laursen, G. Meyn, E. Rasmussen, Toward understanding insulin fibrillation. J Pharm Sci 86, 517–525 (1997).

22. W. Dzwolak et al., Conformational indeterminism in protein misfolding: chiral amplification on amyloidogenic pathway of insulin. J Am Chem Soc 129, 7517–7522 (2007).

23. M. Groenning, S. Frokjaer, B. Vestergaard, Formation mechanism of insulin fibrils and structural aspects of the insulin fibrillation process. Curr Protein Pept Sci 10, 509–528 (2009).

24. E. Chatani et al., Early aggregation preceding the nucleation of insulin amyloid fibrils as monitored by small angle X-ray scattering. Sci Rep 5, 15485 (2015).

25. M. V. Smoluchowski, Drei Vortrage Uber Diffusion, Brownsche Bewegung Und Koagulation Von Kolloidteilchen. Phys Z 17, 557−585 (1916).

26. H. Imamura, S. Honda, Kinetics of Antibody Aggregation at Neutral pH and Ambient Temperatures Triggered by Temporal Exposure to Acid. J Phys Chem B 120, 9581–9589 (2016).

27. R. R. Porter, Partition chromatography of insulin and other proteins. Biochem J 53, 320–328 (1953).

28. B. J. Frisken, Revisiting the method of cumulants for the analysis of dynamic light-scattering data. Appl Opt 40, 4087–4091 (2001).

29. B. J. Berne, R. Pecora, Dynamic Light Scattering: with Applications to Chemistry, Biology, and Physics, John Wiley & Sons, Inc.: New York (1976).

30. C. S. Johnson, D. A. Gabriel, Laser Light Scattering, Dover Publications, Inc.: New York (1994).

31. K. Yuzu et al., Multistep Changes in Amyloid Structure Induced by Cross-Seeding on a Rugged Energy Landscape. Biophys J 120, 284–295 (2021).

32. L. R. De Young, A. L. Fink, K. A. Dill, Aggregation of Globular Proteins. Acc Chem Res 26, 614−620 (1993).

33. T. A. Witten, Jr., L. M. Sander, Diffusion-Limited Aggregation, a Kinetic Critical Phenomenon. Phys Rev Lett 47, 1400−1403 (1981).

34. T. Vicsek, Fractal Growth Phenomena, 2nd ed. World Scientific Pub Co. Inc.: Singapore (1992).

35. D. A. Weitz, J. S. Huang, M. Y. Lin, J. Sung, Limits of the fractal dimension for irreversible kinetic aggregation of gold colloids. Phys Rev Lett 54, 1416–1419 (1985).

36. D. A. Weitz, M. Oliveria, Fractal Structures Formed by Kinetic Aggregation of Aqueous Gold Colloids. Phys Rev Lett 52, 1433−1436 (1984).

37. N. Fuchs, Über Die Stabilitat Und Aufladung Der Aerosole. Phys Z 89, 736−743 (1934).

38. J. Feder, T. Jøssang, E. Rosenqvist, Scaling Behavior and Cluster Fractal Dimension Determined by Light Scattering from Aggregating Proteins. Phys Rev Lett 53, 1403−1406 (1984).

39. T. Jøssangg, J. Feder, E. Rosenqvist, Heat Aggregation Kinetics of Human IgG. J Chem Phys 82, 574−589 (1985).

40. M. Owczarz, A. C. Motta, M. Morbidelli, P. Arosio, A Colloidal Description of Intermolecular Interactions Driving Fibril-Fibril Aggregation of a Model Amphiphilic Peptide. Langmuir 31, 7590–7600 (2015).

41. T. L. Hill, Length dependence of rate constants for end-to-end association and dissociation of equilibrium linear aggregates. Biophys J 44, 285–288 (1983).

42. S. J. Tomski, R. M. Murphy, Kinetics of aggregation of synthetic beta-amyloid peptide. Arch Biochem Biophys 294, 630–638 (1992).

43. M. M. Pallitto, R. M. Murphy, A mathematical model of the kinetics of beta-amyloid fibril growth from the denatured state. Biophys J 81, 1805–1822 (2001).

44. A. J. Modler, K. Gast, G. Lutsch, G. Damaschun, Assembly of amyloid protofibrils via critical oligomers--a novel pathway of amyloid formation. J Mol Biol 325, 135–148 (2003).

45. R. Carrotta, M. Manno, D. Bulone, V. Martorana, P. L. San Biagio, Protofibril formation of amyloid beta-protein at low pH via a non-cooperative elongation mechanism. J Biol Chem 280, 30001–30008 (2005).

46. P. Arosio, M. Owczarz, H. Wu, A. Butte, M. Morbidelli, End-to-end self-assembly of RADA 16-I nanofibrils in aqueous solutions. Biophys J 102, 1617–1626 (2012).

47. V. Fodera, A. Zaccone, M. Lattuada, A. M. Donald, Electrostatics controls the formation of amyloid superstructures in protein aggregation. Phys Rev Lett 111, 108105 (2013).

48. A. J. Dear et al., Kinetic diversity of amyloid oligomers. Proc Natl Acad Sci U S A 117, 12087–12094 (2020).

49. T. C. T. Michaels, A. J. Dear, S. I. A. Cohen, M. Vendruscolo, T. P. J. Knowles, Kinetic profiling of therapeutic strategies for inhibiting the formation of amyloid oligomers. Journal of Chemical Physics 156 (2022).

50. F. Hofmeister, Zur Lehre von der Wirkung der Salze. Arch Exp Pathol Pharmakol 24, 247−260 (1888).

51. T. Kimura et al., Dehydration of main-chain amides in the final folding step of single-chain monellin revealed by time-resolved infrared spectroscopy. P Natl Acad Sci USA 105, 13391–13396 (2008).

52. G. Reddy, J. E. Straubb, D. Thirumalai, Dynamics of locking of peptides onto growing amyloid fibrils. P Natl Acad Sci USA 106, 11948–11953 (2009).

53. D. Thirumalai, G. Reddy, J. E. Straub, Role of Water in Protein Aggregation and Amyloid Polymorphism. Accounts Chem Res 45, 83–92 (2012).

54. C. Nilsberth et al., The ‘Arctic’ APP mutation (E693G) causes Alzheimer’s disease by enhanced Abeta protofibril formation. Nat Neurosci 4, 887–893 (2001).

55. T. Tomiyama et al., A new amyloid beta variant favoring oligomerization in Alzheimer’s-type dementia. Ann Neurol 63, 377–387 (2008).

56. K. Ono, T. Ikeda, J. Takasaki, M. Yamada, Familial Parkinson disease mutations influence alpha-synuclein assembly. Neurobiol Dis 43, 715–724 (2011).

57. S. J. Hamodrakas, Protein aggregation and amyloid fibril formation prediction software from primary sequence: towards controlling the formation of bacterial inclusion bodies. FEBS J 278, 2428–2435 (2011).

