## Supplementary material for "Mechanistic modeling of amyloid oligomer and protofibril formation": SI

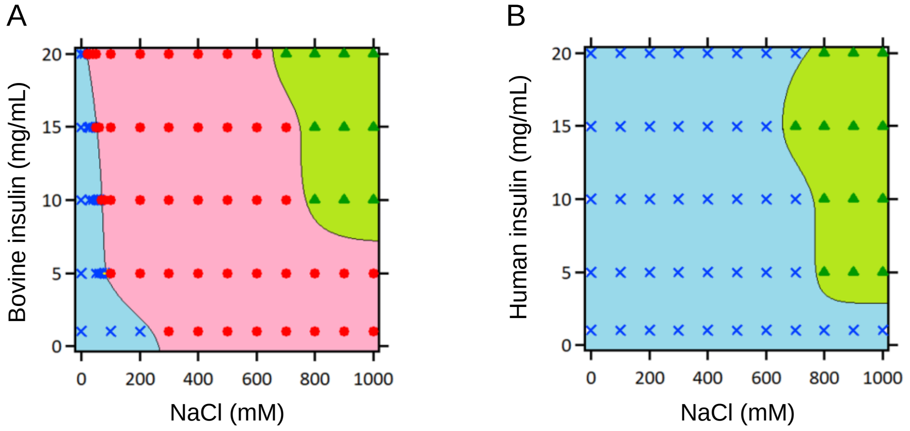


**Figure S1.** Phase diagram of amyloid fibril formation pathway of bovine (A) and human insulin (B). The blue and red areas indicate amyloid fibril formation pathway without and with prefibrillar aggregates, respectively. The green area indicates amorphous aggregation pathway. The phase diagrams were constructed based on the results of ThT fluorescence assay.

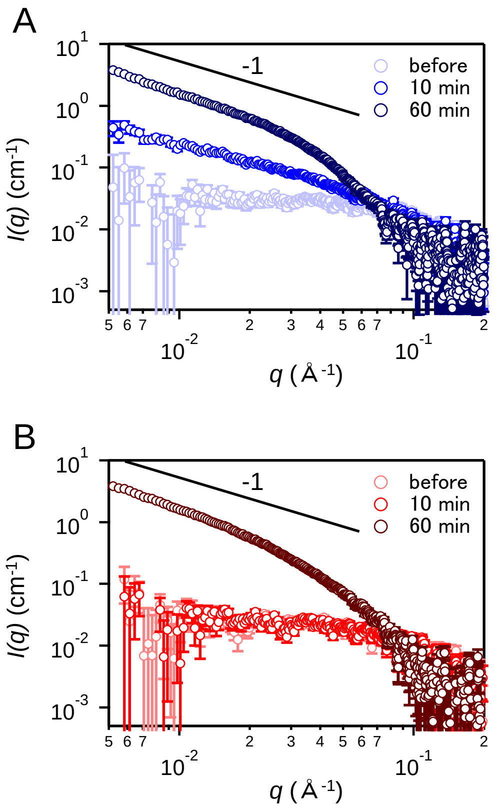


**Figure S2.** SAXS profiles of aggregated species of bovine (A) and human insulin (B). Samples of 5.0 mg/mL insulin dissolved in 25 mM HCl containing 0.1 M NaCl was heated at 75 °C for 10 and 60 min, and the reaction was instantly stopped with cooling on ice. SAXS measurements were conducted with a laboratory-based instrument NANOPIX (Rigaku, Japan). The sample-to-detector distance (SDD) was set to be 1330 mm, with which the covered *q*-range was 0.005 $Å^{-1} \leq q \leq 0.20$ $Å^{-1}$. All measurements were performed at 25 °C. One-dimensional scattering profiles $I(q)$ were obtained by the radial averaging of two-dimensional scattering patterns using SAngler software (1). After correction by the transmittance and subtraction of buffer scattering, the absolute scattering intensity was obtained using the standard scattering intensity of water ($1.632 \times 10^{-2}$ $cm^{-1}$) (2).

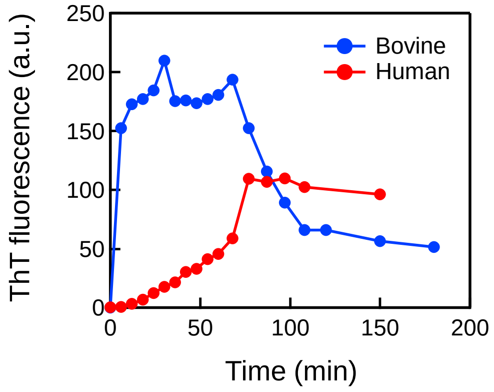


**Figure S3.** Time courses of ThT fluorescence intensity of amyloid fibril formation of bovine and human insulin under deuterated conditions. Prior to the sample preparation, bovine insulin was preincubated in 25 mM DCl and at 37 °C for 24 h to complete the H/D exchange of amide protons, and 5.0 mg/mL of insulin solution dissolved in 25 mM DCl containing 0.5 M NaCl was heated at 75 °C. At different time points, a 4.5 μL aliquot of the sample solution was mixed with 1.5 mL of 5 μM ThT in 50 mM Gly-NaOH buffer (pH 8.5), and the fluorescence intensity at 485 nm was measured using an excitation wavelength of 445 nm with a RF-5300 PC spectrofluorometer (Shimadzu Corporation, Kyoto, Japan).

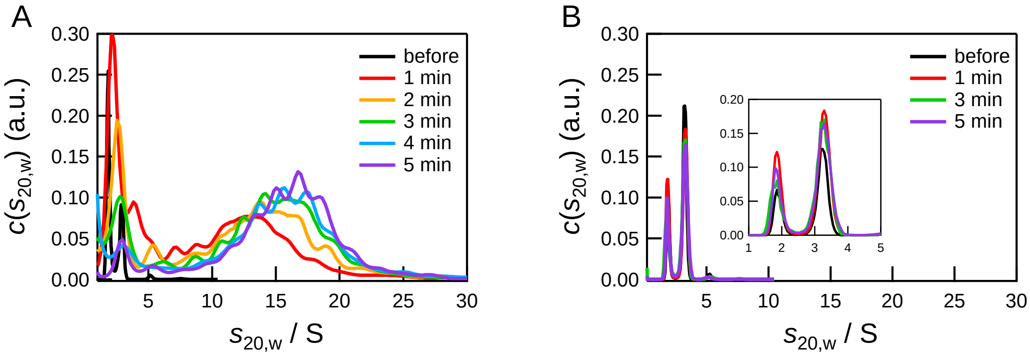


**Figure S4.** Sedimentation coefficient distributions of early aggregated species of bovine (A) and human insulin (B). Analytical ultracentrifugation (AUC) was conducted with a ProteomeLab XL-I analytical ultracentrifuge (Beckman Coulter, Brea, CA, USA). A sample of 5.0 mg/mL insulin dissolved in 25 mM HCl containing 0.5 M NaCl was heated at 75 °C for different times ranging from 1 to 5 min, and the reaction was instantly stopped with cooling on ice. For bovine insulin, sample solutions were diluted 2-fold with 25 mM HCl to suppress further flocculation of aggregates induced by cooling. Sedimentation velocity analytical ultracentrifugation (SV-AUC) measurements were performed using Rayleigh interference optics at 40,000 rpm at 20 °C with a 1.5 mm path-length cell. The experimental data of SV-AUC were analyzed with SEDFIT software which executed the fitting with Lamm formula (3). For the analysis, the density and viscosity of solvents, and partial specific volumes of each protein were calculated from their amino acid sequences with SEDNTERP software (4).

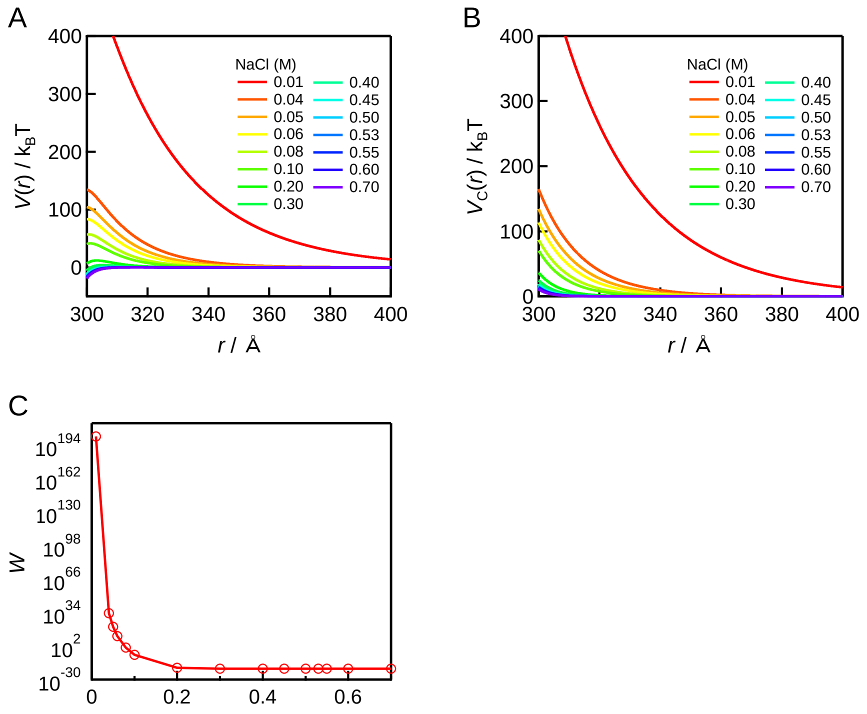


**Figure S5.** Salt concentration dependence of intermolecular potential of a protein molecule, $V(r)$ (A), screened repulsive Coulomb potential, $V_C(r)$ (B), and Fuchs stability ratio, $W$ (C). Based on the Derjaguin-Laudau-Verwey-Overbeek (DLVO) theory, $V(r)$ is expressed as

$$V(r) = V_{\mathrm{C}}(r) + V_{\mathrm{AY}}(r)$$

where $r$ is the distance between the protein molecules. $\sigma$ is the diameter of the protein. The terms, $V_C(r)$ and $V_{AY}(r)$ are the screened repulsive Coulomb potential and the attractive potential, respectively. Here, $V_{AY}(r)$ is assumed to be Yukawa-type potential:

$$V_{\mathrm{C}}(r) = \frac{Z^2 e^2}{\varepsilon(1+0.5\kappa\sigma)^2} \frac{\exp[-\kappa(r-\sigma)]}{r}$$

$$V_{\mathrm{AY}}(r) = -J\left(\frac{\sigma}{r}\right)\exp\left[\frac{-(r-\sigma)}{d}\right]$$

$Z$ is the net charge on the protein, $e$ is the elementary charge, $\varepsilon$ is the dielectric constant of the medium, $\kappa$ is the reciprocal Debye-Hückel screening length, $J$ is the depth of attractive potential at $r = s$, and $d$ is the range of the attractive potential. In the present calculation, we used $J = 30\ k_B T$, $d = 3$ Å, $Z = +900$ (+6×150 mer), and $\sigma = 300$ Å (30 nm).

**Table S1.** Effective aggregation rate constant for the temperature dependence on early aggregation of bovine insulin obtained by fitting analysis

| Temperature (℃) | threshold association number for the lag phase in the first step | threshold association number for the second step | $k_{eff}$ ($m^3$ $sec^{-1}$) | | |
|---|---|---|---|---|---|
| | | | first step (lag phase) | first step (growth phase) | second step |
| 70 | 30 | 135 | $2.7 (\pm 0.5) \times 10^{-25}$ | $1.2 (\pm 0.8) \times 10^{-23}$ | $4.1 (\pm 3.7) \times 10^{-27}$ |
| 72 | 30 | 145 | $3.4 (\pm 0.3) \times 10^{-25}$ | $1.2 (\pm 0.3) \times 10^{-23}$ | $5.1 (\pm 2.0) \times 10^{-27}$ |
| 75 | 30 | 150 | $5.2 (\pm 0.2) \times 10^{-25}$ | $9.0 (\pm 0.5) \times 10^{-24}$ | $1.4 (\pm 0.1) \times 10^{-26}$ |
| 78 | 30 | 160 | $7.0 (\pm 0.2) \times 10^{-25}$ | $1.0 (\pm 0.04) \times 10^{-23}$ | $1.6 (\pm 0.1) \times 10^{-26}$ |
| 80 | 30 | 170 | $1.0 (\pm 0.04) \times 10^{-24}$ | $1.0 (\pm 0.03) \times 10^{-23}$ | $2.3 (\pm 0.1) \times 10^{-26}$ |

**Table S2.** Effective aggregation rate constant for the salt concentration dependence on early aggregation of bovine insulin obtained by fitting analysis

| NaCl (M) | threshold association number for the lag phase in the first step | threshold association number for the second step | $k_{eff}$ ($m^3$ $sec^{-1}$) | | |
|---|---|---|---|---|---|
| | | | first step (lag phase) | first step (growth phase) | second step |
| 0.40 | 30 | 135 | $5.5\ (\pm 0.4) \times 10^{-25}$ | $7.5\ (\pm 0.7) \times 10^{-24}$ | $5.5\ (\pm 2.1) \times 10^{-27}$ |
| 0.45 | 30 | 145 | $5.8\ (\pm 0.3) \times 10^{-25}$ | $8.0\ (\pm 0.5) \times 10^{-24}$ | $1.2\ (\pm 0.2) \times 10^{-26}$ |
| 0.50 | 30 | 150 | $5.5\ (\pm 0.2) \times 10^{-25}$ | $8.2\ (\pm 0.4) \times 10^{-24}$ | $1.5\ (\pm 0.1) \times 10^{-26}$ |
| 0.53 | 30 | 160 | $4.8\ (\pm 0.2) \times 10^{-25}$ | $9.9\ (\pm 0.6) \times 10^{-24}$ | $1.4\ (\pm 0.2) \times 10^{-26}$ |
| 0.55 | 30 | 170 | $5.0\ (\pm 0.2) \times 10^{-24}$ | $9.1\ (\pm 0.5) \times 10^{-23}$ | $2.0\ (\pm 0.2) \times 10^{-26}$ |

**Table S3.** Effective aggregation rate constant for the Hofmeister salt dependence on early aggregation of bovine insulin obtained by fitting analysis.

| Salt | threshold association number for the lag phase in the first step | threshold association number for the second step | $k_{eff}$ ($m^3\ sec^{-1}$) | | |
|---|---|---|---|---|---|
| | | | first step (lag phase) | first step (growth phase) | second step |
| LiCl | 30 | 145 | $5.5\ (\pm 0.2) \times 10^{-25}$ | $9.1\ (\pm 0.6) \times 10^{-24}$ | $1.2\ (\pm 0.2) \times 10^{-26}$ |
| NaCl | 30 | 145 | $5.8\ (\pm 0.3) \times 10^{-25}$ | $8.0\ (\pm 0.5) \times 10^{-24}$ | $1.2\ (\pm 0.2) \times 10^{-26}$ |
| KCl | 30 | 140 | $4.3\ (\pm 0.3) \times 10^{-25}$ | $1.1\ (\pm 0.1) \times 10^{-23}$ | $9.6\ (\pm 1.9) \times 10^{-27}$ |
| RbCl | 30 | 130 | $4.4\ (\pm 0.3) \times 10^{-25}$ | $1.3\ (\pm 0.2) \times 10^{-23}$ | $7.4\ (\pm 1.9) \times 10^{-27}$ |
| CsCl | 30 | 130 | $4.2\ (\pm 0.4) \times 10^{-25}$ | $1.2\ (\pm 0.3) \times 10^{-23}$ | $3.8\ (\pm 2.1) \times 10^{-27}$ |
| $NH_4Cl$ | 30 | 125 | $4.6\ (\pm 0.4) \times 10^{-25}$ | $1.2\ (\pm 0.2) \times 10^{-23}$ | $8.5\ (\pm 2.2) \times 10^{-27}$ |

### References


1. N. Shimizu *et al.*, Software Development for Analysis of Small-angle X-ray Scattering Data. *Aip Conf Proc* **1741** (2016).
2. D. Orthaber, A. Bergmann, O. Glatter, SAXS experiments on absolute scale with Kratky systems using water as a secondary standard. *J Appl Crystallogr* **33**, 218-225 (2000).
3. P. Schuck, Size-distribution analysis of macromolecules by sedimentation velocity ultracentrifugation and Lamm equation modeling. *Biophys J* **78**, 1606-1619 (2000).
4. J. S. Philo, SEDNTERP: a calculation and database utility to aid interpretation of analytical ultracentrifugation and light scattering data. *Eur Biophys J Biophy* 10.1007/s00249-023-01629-0 (2023).
